# A Visuomotor Model Enhances Human Position Sense

**DOI:** 10.1101/2020.07.20.211490

**Authors:** Charitha Omprakash, Seyedsina Razavizadeh, Max-Philipp Stenner

## Abstract

Integrating information from multiple sources reduces uncertainty. Besides sensory input, animals have access to another source of information about their body and the environment, i.e., their own motor commands, which alter the body and environment in a predictable way. Does this predictability reduce perceptual uncertainty, i.e., variance? Participants moved their unseen arm and reported movement endpoint locations. In two conditions, a predictive model of visuomotor contingencies could either be fully formed, and used for this estimation, or remained incomplete. This was achieved through context trials that provided visual endpoint feedback at a predictable vs. unpredictable latency, while carrying identical spatial information. In two experiments, we found that endpoint estimation was less variable when a full, spatiotemporal, model could be formed. Higher perceptual precision was paralleled by enhanced movement accuracy. We conclude that a visuomotor model provides a separate source of information, additional to sensory input, which enhances human position sense.

## Introduction

Integrating information from multiple sources can reduce uncertainty and variance (Ernst & Banks, 2002). Besides sensory input, animals have access to another key source of information about the body and the environment, that is, their own motor commands. Voluntary movement alters the state of the body and the environment in a controlled, i.e., predictable, way. Does this predictability reduce uncertainty, and, thus, variance, in perceiving movement-induced state changes?

There is ample evidence that the nervous system estimates how imminent motor output will change the state of the body (e.g., Duhamel, Colby, & Goldberg, 1992; Guthrie, Porter, & Sparks, 1983; Wolpert, Ghahramani, & Jordan, 1995; for a review see Shadmehr, Smith, & Krakauer, 2010). These estimates, thought to be derived from an internal model of sensorimotor contingencies, are considered critical for accurate, flexible motor control (e.g., Franklin & Wolpert, 2011; Shadmehr et al., 2010). There is also evidence that internal model estimates can systematically bias perception (e.g., Bays, Wolpert, & Flanagan, 2005; Izawa, Criscimagna-Hemminger, & Shadmehr, 2012; Synofzik, Lindner, & Thier, 2008). However, it is largely unclear how predictability derived from motor outflow influences perceptual uncertainty. This is important because inverse uncertainty – precision – is a hallmark of reliable perception (Ernst & Banks, 2002; Ernst & Bülthoff, 2004).

Few studies have investigated whether, and how, predictability of the consequences of motor outflow changes perceptual precision. This is surprising given recent evidence that predictability in a broader sense, derived from statistical regularities in the environment, can indeed “sharpen” sensory representations (Kok, Jehee, & de Lange, 2012; Summerfield & Lange, 2014). The few previous investigations into perceptual precision in the context of movement have remained difficult to interpret due to ambiguities raised by their designs. Specifically, previous studies have changed sensory feedback in a task-relevant dimension to operationalise predictability (Bhanpuri, Okamura, & Bastian, 2013; Vaziri, Diedrichsen, & Shadmehr, 2006), or introduced important potential confounds of predictability with conflict across sensory modalities (Yon et al., 2018; see discussion for more details).

Here, we avoid typical ambiguities in previous studies. We focus on human position sense. Position sense profits from multisensory integration of proprioception and vision (van Beers, Sittig, & Denier van der Gon, 1996; van Beers, Wolpert, & Haggard, 2002). However, movements often unfold in the absence of visual feedback, e.g., when reaching out to shift gears while steering a car. Do humans purely rely on proprioception in these cases? Or do we enhance precision by taking into account how motor outflow will change the state of the body? This is important because reliable, precise estimation of limb position guides movement planning and correction (Desmurget, Pelisson, Rossetti, & Prablanc, 1998; Desmurget, Rossetti, Jordan, Meckler, & Prablanc, 1997; but see Polit & Bizzi, 1978).

To address this question, we examine spatial precision of localizing the unseen arm after a reaching movement in two contexts of high vs. low predictability of movement feedback. Importantly, to avoid typical ambiguities in previous studies, we manipulate predictability along a task-irrelevant sensory dimension – time – and hold spatial information constant across conditions. Previous evidence suggests that internal models are tuned both in space and time (Bays et al., 2005; Blakemore, Frith, & Wolpert, 1999; Kilteni, Houborg, & Ehrsson, 2019). Accordingly, we expected that a disruption of temporal predictability of movement consequences would also impair their internal spatial estimation. If this internal spatial estimation contributes to position sense, the latter should thus become less precise, i.e., more variable, when temporal predictability decreases.

We indeed provide evidence that human position sense is more precise when a full, spatiotemporal, visuomotor model can be formed. We discuss potential mechanisms of this increase in precision, together with strong potential implications for an interaction of different motor learning mechanisms (e.g., Mazzoni & Krakauer, 2006).

## Methods

### Participants

23 right-handed participants took part in Experiment 1 (10 female; age (mean ± SD): 25.7 ± 1.4 yrs; Edinburgh handedness inventory (Oldfield, 1971) (mean ± SD): 82.7 ± 9.3). 22 right-handed participants took part in Experiment 2 (11 female; age (mean ± SD): 25.7 ±1.6 yrs; Edinburgh handedness inventory (mean ± SD): 89.5 ± 6.6). Sample sizes for both experiments were at least twice as large as previous psychophysical studies into effects of internal models on perceptual precision (Bhanpuri et al., 2013; Vaziri et al., 2006). All participants were recruited from staff and students at Otto-von-Guericke University. All had normal or corrected-to-normal vision, and were free from any known neurological disorder. The study was approved by the local ethics committee at Otto-von-Guericke University Magdeburg. Participants gave written informed consent. They were reimbursed for participation (8 € per hour).

### Apparatus

Experiments were conducted in a dimly lit room. Participants were sitting at an experimental table with three levels (**Figure 1A**). The lowermost level held a graphics tablet (Intuos 4XL, Wacom, Kazo, Japan; 5080 lines per inch, sampled at 200 Hz) with an active area of 48.8 x 30.5 cm. The uppermost level held an LCD monitor (52 x 32.4 cm, refresh rate 60 Hz) facing downwards. Participants looked down into a mirror placed at the middle level, equidistant between the monitor and the graphics tablet. The virtual image of the monitor thus appeared in the same plane as the tablet. During the experiment, participants moved a stylus held in their right hand across the graphics tablet and saw visual feedback of their movement endpoint location in the mirror. Their hands, arms and shoulders were hidden from sight beneath a black apron attached to the middle level of the experimental table. Their left hand was resting on a trackball (Experiment 1) or a keypad with five keys arranged in a cross (Experiment 2) placed to the left of the graphics tablet. Throughout the experiment, participants wore polarizing sunglasses to minimize visibility of a second, slightly offset reflection of the monitor at the glass surface of the mirror. They also wore headphones for auditory feedback.

**Figure 1.**
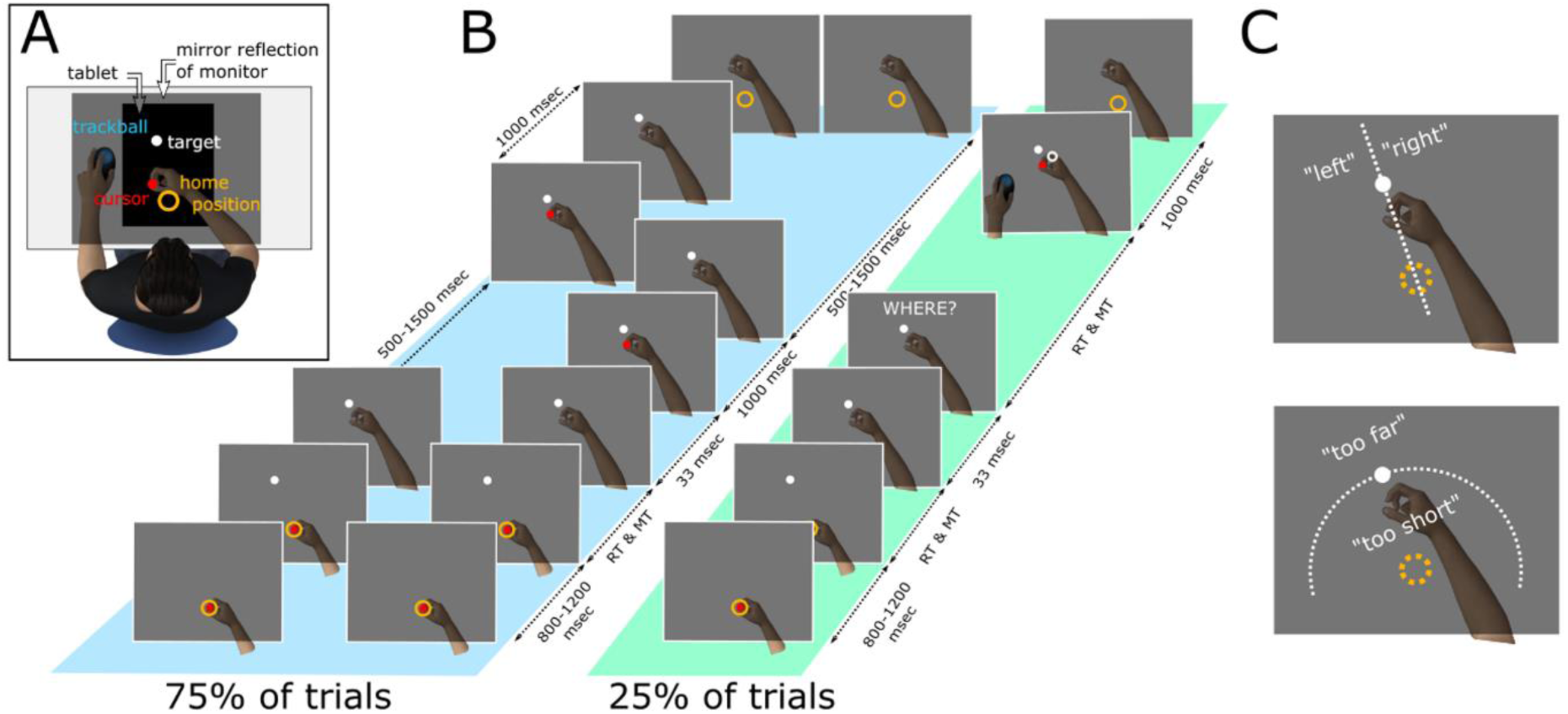
Schematic of the experimental setup and behavioural task. *A*, Experimental setup. For illustrative purposes, the middle level of the experimental table is depicted as transparent, while it actually occluded arms, hands, and the tablet from sight. In addition, participants were wearing an apron so that shoulders were not visible either. *B*, Schematic of exemplary trials across different conditions. Blue, context trials. Left column, context trial in the unpredictable condition. Middle column, context trial in the predictable condition. Right column (green), perceptual trial in Experiment 1. The word “Where” appeared at the bottom of the screen but is here shown at the top for readability. In Experiment 2, instead of estimating movement endpoint location using a trackball, participants made two consecutive two-alternative forced choices, indicating whether their movement endpoint was “left” or “right” and “too far” or “too short” relative to the target location. *C*, Definition of response categories in the two-alternative forced choice task in Experiment 2. The text in *C*, and all dashed lines, were never presented on the screen, and are shown purely for illustrative purposes. Schematics of the visual display and of visual stimuli in *A, B*, and *C* are not to scale.

### Behavioral Tasks

The behavioral tasks were programmed and presented using Presentation (Neurobehavioral Systems). In both experiments, participants were asked to make rapid, accurate reaching movements with their unseen right hand to visual targets displayed in the mirror. Following some of these movements, participants estimated movement endpoints in the absence of visual feedback. In the remaining trials, veridical visual feedback of movement endpoints was provided, either immediately after movement offset (predictable condition), or with a varying temporal delay (unpredictable condition, tested in a separate experimental session on a different day). The goal of the behavioral task was to examine how spatial precision in locating unseen movement endpoints changed when a (temporal) prediction of visual feedback could be formed (predictable vs. unpredictable condition).

In both predictable and unpredictable conditions, each trial started with the presentation of a visual home position, consisting of a yellow circle (0.6 cm diameter, 0.2 cm line width) at the tablet midline, roughly aligned with the body midline (**Figure 1B**). Upon seeing the home position, participants moved the stylus towards the location of the home position. Once the stylus was within 2 cm of the center of the home position, participants saw a red dot (0.4 cm diameter) that indicated the stylus’ current location. After reaching the home position and holding the stylus in place for a random interval between 800 and 1200 ms (uniform distribution) a visual target was presented. The target consisted of a white dot (0.3 cm diameter). To avoid stereotyped movements, target location varied slightly from trial to trial, with targets placed at a distance between 14.5 and 15.5 cm from the home position (uniform distribution), and at an angle between ± 1.9° (uniform distribution) relative to the tablet midline. Participants had to make a straight, rapid reaching movement to bring the stylus as accurately as possible to the perceived location of the target. The time of movement initiation was not restricted. Once the stylus left the home position, both the home position and the red dot disappeared, i.e., no online feedback of the movement was available, while the target remained visible. Movement offset was defined as the time when no further change in location of the stylus was detectable for at least 33 ms. Movement endpoint location was defined as the stylus location at movement offset.

There were two types of trials, context trials and perceptual trials, which differed in the sequence of events following movement offset. Context and perceptual trials were pseudo-randomly interleaved.

75% of trials were context trials (**Figure 1B**, blue). In context trials, participants received visual feedback of their movement endpoint at a latency relative to movement offset that depended on the experimental condition. In both conditions, visual endpoint feedback consisted of the red dot reappearing at the location of movement endpoint for 1000 ms. In the predictable condition, the red dot reappeared immediately after movement offset. In the unpredictable condition, there was a random delay between 500 and 1500 ms between movement offset and reappearance of the red dot (uniform distribution, in steps of 16.67 ms). The two conditions were blocked, with each condition tested on a separate day, a week apart.

If a movement endpoint was within 0.4 cm of the center of the target, there was a visual display of “target explosion”, and participants heard a success sound (a bell sound). Both the explosion and the sound were presented immediately after movement offset in the predictable condition, and with the above random delay of 500 – 1500 ms in the unpredictable condition. Target explosion and success sound were intended to motivate participants to produce accurate, targeted movements.

Following movement offset participants had to hold the stylus in place until the home position reappeared. In the unpredictable condition, the home position reappeared once the red dot indicating movement endpoint location disappeared. In the predictable condition, participants had to wait for a varying time interval after disappearance of the red dot before the home position reappeared. These intervals were chosen such that delays between movement offset and reappearance of the home position were matched between predictable and unpredictable conditions (**Figure 1B**, blue).

The remaining 25% of trials were perceptual trials (**Figure 1B**, green). In perceptual trials, participants did not receive visual feedback of their movement endpoint until they estimated its location. The method participants used to report estimated movement endpoints differed between Experiment 1 and Experiment 2.

In Experiment 1, participants saw the word “Where?” at the bottom of the mirror. This instructed them to move a trackball with their left hand in order to move a visual “copy” of the target, a white ring (0.4 mm diameter, line width 0.2 mm), from the target location, where it was first presented, to the estimated movement endpoint. Once the white ring was at the estimated movement endpoint, participants pressed a key on the trackball. The red dot then immediately reappeared at the actual movement endpoint location. If the white ring was within 0.4 cm of the actual movement endpoint location, the target exploded immediately and the success sound was presented. Perceptual trials were thus experienced, and introduced in the instructions to the task, as a “second chance” to make the target explode.

Experiment 2 tested whether results from Experiment 1 depended on the method participants used to report estimated movement endpoints. Experiment 2 was designed as a two-alternative forced choice task. Participants saw two questions at the bottom of the mirror, one presented after the other. The first question asked “Left or right?”, prompting participants to indicate whether their movement endpoint location was left or right of a straight line connection between the home position and the target (**Figure 1C**, upper panel). To respond, they used left and right keys on a keypad with their left index finger. Upon responding, they received feedback whether their response was correct or not. They saw a green tick next to the question if their response was correct, and a red cross if it was incorrect. This was followed by a second question asking “Too far or too short?”, prompting participants to indicate whether they over- or undershot the target (**Figure 1C**, lower panel). Participants responded by pressing upper or lower keys on the keypad, respectively, again receiving feedback. Following the second response, they saw the red dot reappear at the location of their movement endpoint.

Across all trials, and in both experiments, there was a minimum required average movement speed (20 cm/sec; radial extent of movement divided by movement time, from leaving the home position to movement offset). If participants moved too slowly, they heard an unpleasant error sound upon movement offset (superimposed 600 and 700 Hz sine waves, 200 ms duration). Across all trials and both experiments, participants were also asked to hold the stylus in place following movement offset until the home position reappeared. If they moved the stylus by more than 1 mm during this time, another unpleasant error sound was played (superimposed 1000 and 1200 Hz sine waves, 100 ms duration). This ensured that participants were not moving and generating additional proprioceptive feedback, which could have been otherwise used for movement endpoint estimation. Upon reappearance of the home position, participants returned to the home position to start the next trial.

### Experimental Procedure

Each condition, predictable and unpredictable, was completed on a separate day, a week apart. The order of conditions was counterbalanced across subjects (the predictable condition was completed first by 12 participants in Experiment 1, and by 14 participants in Experiment 2). Each session lasted for approximately two hours on each day, including instructions and training. Participants first trained to execute rapid and accurate reaching movements to visual targets. They then trained to additionally estimate movement endpoints. The actual experiment consisted of eight blocks of 52 trials each. Between blocks, participants saw a performance score on the screen, which reflected how close their movements were, on average, to the target.

### Analysis

Data from both experiments were analysed in MATLAB 2019a and R 3.6.2. We excluded one participant from Experiment 2 due to an excessive number of movements that did not fulfil the predefined criterion for minimum movement speed, and where thus followed by an error sound (see *Behavioral Tasks*; 275 of a total of 416 movements were too slow). For the remaining participants, we excluded trials based on four criteria. We excluded all trials with an error sound due to slow movement speed (on average 35 trials in Experiment 1 and 16 trials in Experiment 2). We also excluded trials with large deviations of movement endpoints from targets (>20° error in direction, or >6 cm error in extent), and movements whose trajectories were strongly curved (linearity index >2, defined as the maximum perpendicular distance of a movement trajectory to a straight line connection between home position and endpoint, divided by the distance between home position and endpoint; Atkeson & Hollerbach, 1985). Based on these criteria, on average four trials were excluded in Experiment 1, and six trials, on average, in Experiment 2. Finally, to minimize any influence of renewed proprioceptive input following movement offset, we excluded trials in which the stylus moved by more than 1 mm following movement offset (resulting in an error sound) from all analyses of estimated movement endpoints (Figures 3 and 5; on average 13 and 12 perceptual trials in Experiments 1 and 2, respectively). On average, kinematic analyses (Figures 2 and 4) were based on 380 trials in Experiment 1 and on 396 trials in Experiment 2. Analyses of estimated endpoints (Figures 3 and 5) were based, on average, on 84 and 90 perceptual trials in Experiments 1 and 2, respectively.

**Figure 2.**
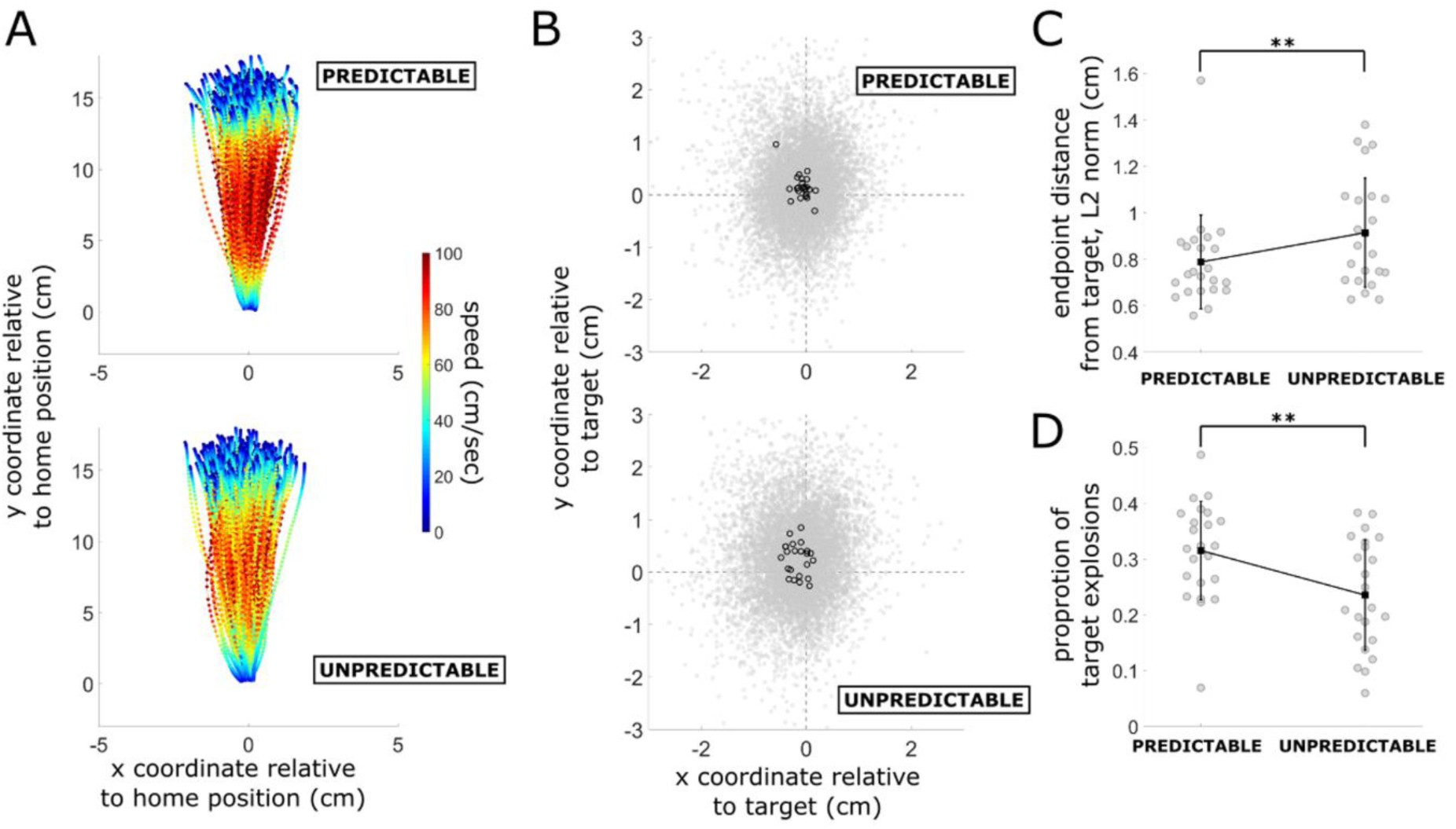
Movement kinematics in Experiment 1. *A*, Spatial trajectories and velocity profiles of all movements across context trials and perceptual trials of a typical participant in the predictable (top) and unpredictable (bottom) condition. *B*, Movement endpoints relative to target location (at (0,0)) across context trials and perceptual trials for all participants in the predictable (top) and unpredictable (bottom) condition. Each black circle represents the mean endpoint location for one participant. *C*, Mean Euclidean distance between movement endpoint and target location across context and perceptual trials for each participant in the predictable (left) and unpredictable (right) condition. *D*, Proportion of target explosions for each participant in the predictable (left) and unpredictable (right) condition. Black rectangles and error bars in *C* and *D* show mean and standard error, respectively. **, p<.01.

**Figure 3.**
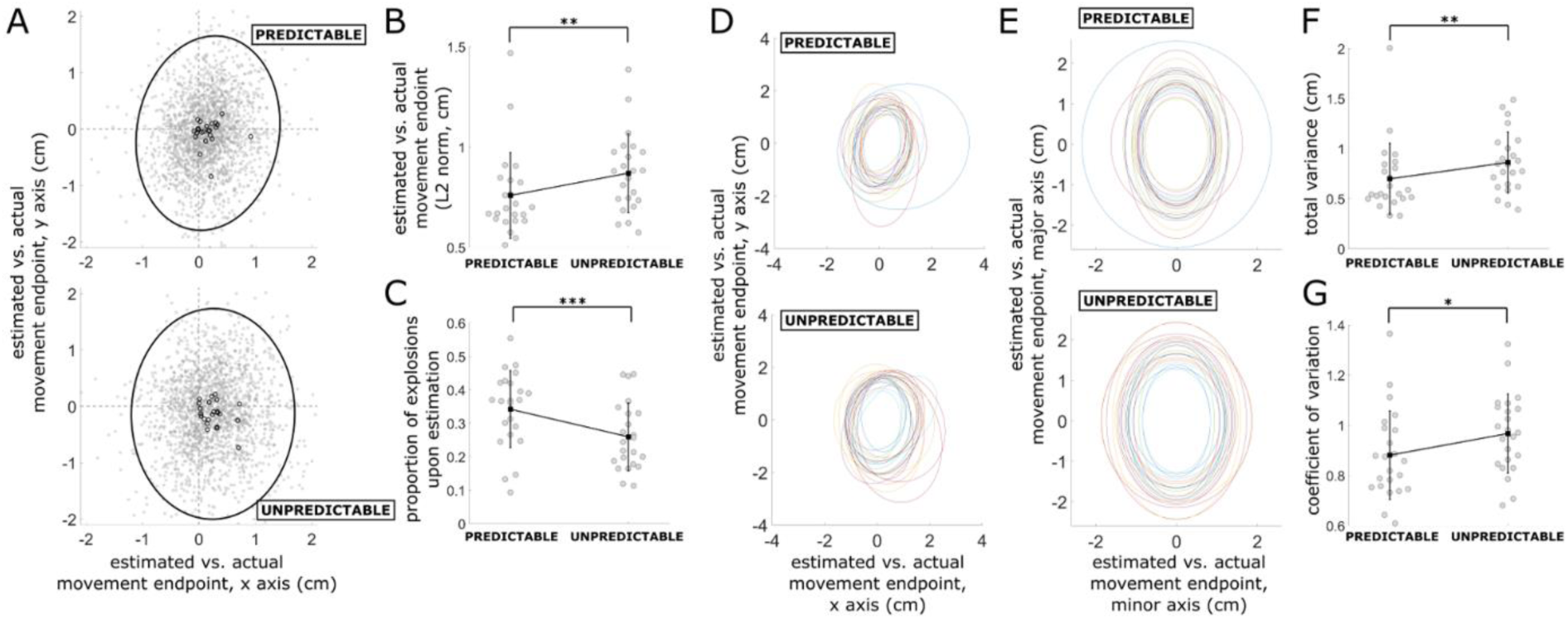
Endpoint estimation precision and accuracy in Experiment 1. *A*, Estimated movement endpoint locations, relative to actual movement endpoint locations (at (0,0)), across all trials and all participants in the predictable (top) and unpredictable (bottom) condition. Each small black circle represents the mean estimated endpoint location for one participant. Ellipses represent 95% confidence regions across all trials and all participants. *B*, Mean Euclidean distance between estimated and actual movement endpoint location for each participant in the predictable (left) and unpredictable (right) condition. *C*, Proportion of target explosions upon endpoint estimation for each participant in the predictable (left) and unpredictable (right) condition. *D*, 95% confidence ellipses for estimated relative to actual movement endpoint locations in the predictable (top) and unpredictable (bottom) condition. Each colour represents a different participant. E, Same as in D, but after removing systematic estimation biases in individual subjects, and rotating estimated endpoint locations such that eigenvectors of individual-subject covariance matrices are aligned. *F*, Total variance (sum of variances in x- and y-dimensions) in the predictable (left) and unpredictable (right) condition. *G*, Coefficient of variation in the predictable (left) and unpredictable (right) condition. Black rectangles and error bars in *B, C, F*, and *G* show mean and standard error, respectively. *, p<.05, **, p<.01, ***, p<.001.

**Figure 4.**
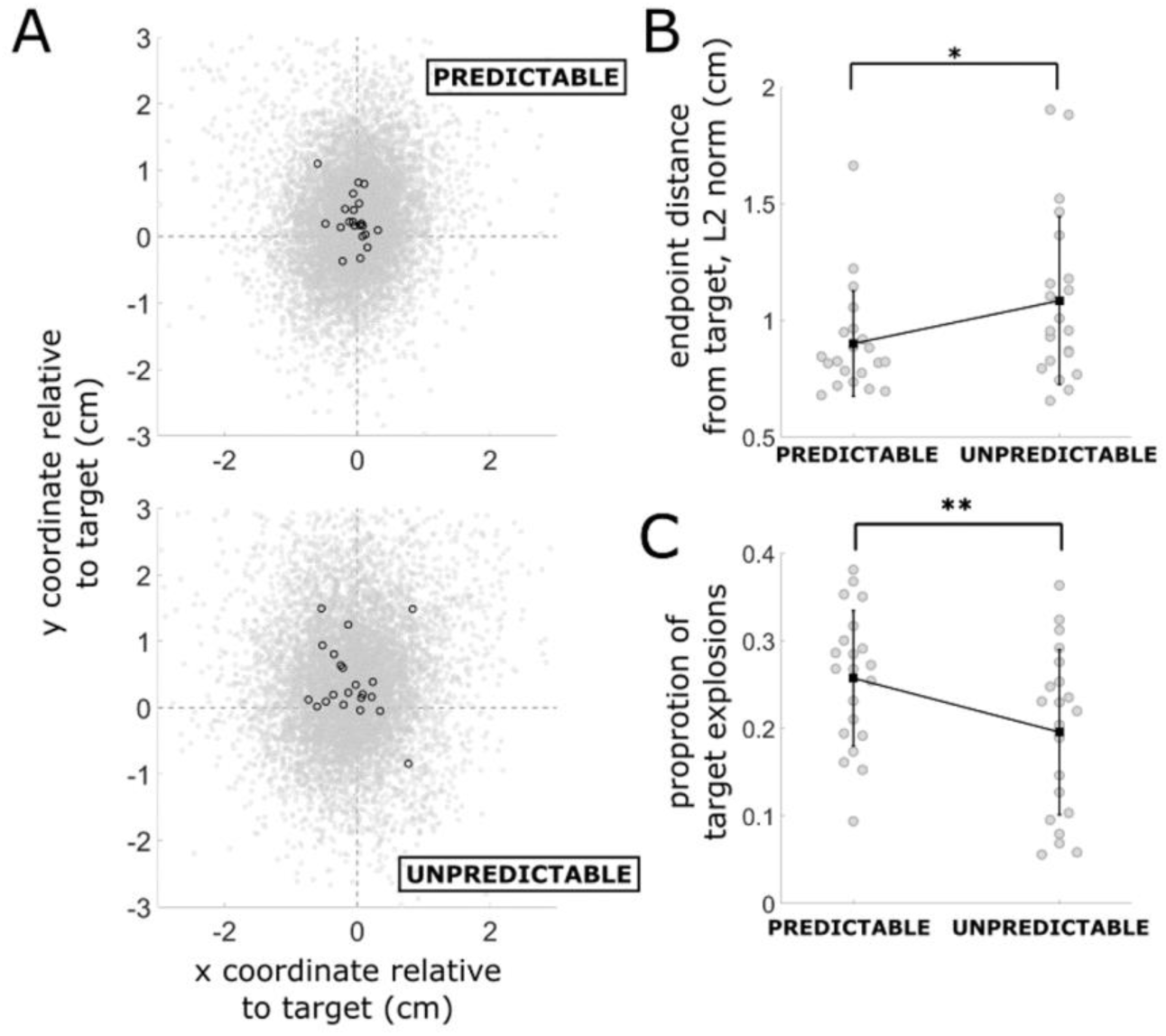
Movement kinematics in Experiment 2. *A*, Movement endpoints relative to target location (at (0,0)) across context trials and perceptual trials for all participants in the predictable (top) and unpredictable (bottom) condition. Each black circle represents the mean endpoint location for one participant. *B*, Mean Euclidean distance between movement endpoint and target location across context and perceptual trials for each participant in the predictable (left) and unpredictable (right) condition. *C*, Proportion of target explosions for each participant in the predictable (left) and unpredictable (right) condition. Black rectangles and error bars in *B* and *C* show mean and standard error, respectively. *, p<.05, **, p<.001.

**Figure 5.**
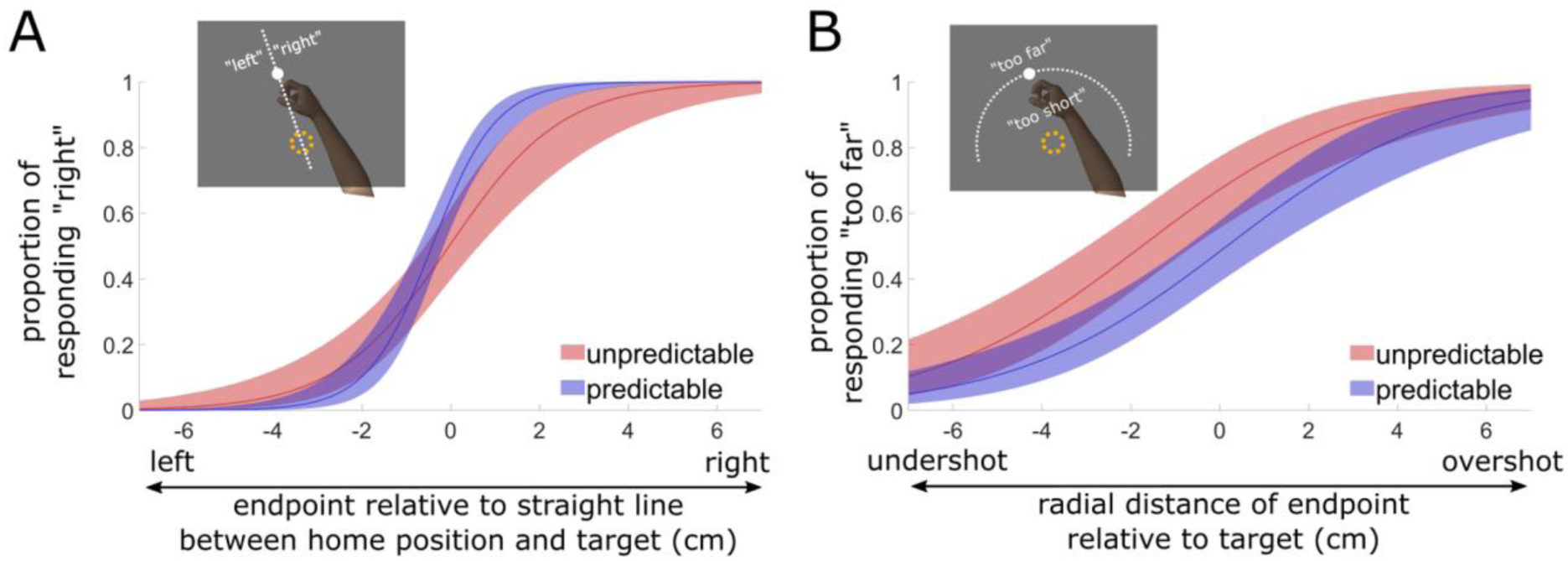
Endpoint estimation precision in Experiment 2. *A*, Marginal predicted values for the proportion of responding “right” (y-axis) as a function of directional movement endpoint error (x-axis) in the predictable (blue) and unpredictable (red) condition. The proportion of responding “right” was predicted from the winning generalized linear mixed-effects model, which included, as fixed-effects, directional movement error, experimental condition (predictable vs. unpredictable), as well as their interaction. Shading represents estimated 95% confidence intervals. Spatial resolution in perceiving movement error is evident in the slope of each sigmoid. The interaction between movement error and condition (the difference in slopes) thus represents differences in spatial resolution, i.e., perceptual precision, between conditions. *B*, Same as A, but for the two-alternative forced choice of responding “too far” or “too short”. Here, the winning generalized linear mixed-effects model included extent error, experimental condition, but not their interaction. Insets in *A* and *B* show schematics of instructed response criteria.

For both experiments we computed movement accuracy, defined as the trial-mean Euclidean distance between movement endpoint location and target location, as well as the proportion of target explosions, and the trial-mean peak movement velocity, computed from the derivative of stylus position after smoothing with a second order Savitzky-Golay filter (frame length of 31 samples, corresponding to 155 ms; e.g., Wong, Marvel, Taylor, & Krakauer, 2019). For Experiment 1, we additionally computed endpoint estimation accuracy, defined as the trial-mean Euclidean distance between estimated and actual movement endpoints, estimation precision, defined as the sum of variances in the (signed) distance between estimated and actual movement endpoints along x and y workspace dimensions (van Beers, 2003), the coefficient of variation (estimation precision divided by estimation accuracy), and the proportion of target explosions following estimation of movement endpoints. These dependent variables were compared between predictable and unpredictable conditions using two-sided dependent-samples t-tests or, when a Shapiro-Wilk test indicated violations of the normality assumption, by a Wilcoxon signed-rank test (using Jasp).

In Experiment 2, estimation precision was computed based on binary responses in the two-alternative forced choice task (“left/right” and “too far/too short”). To this end, we conducted a generalized linear mixed-effects model analysis using the *glmmadaptive* package (Rizopoulos, 2019) in R (R Core Team, 2016). Participants’ binary responses in the two-alternative forced choice task were modelled using mixed-effects logistic regression. Estimation of movement direction (“left/right” choices) and movement extent (“too far/too short” choices) were modelled separately. We expected responses to depend on movement endpoints relative to the target. That is, the more movement endpoints were to the right of a straight line connection between home position and target, the more likely participants were expected to report “right”, and the more movements overshot the target, the more likely participants were expected to report “too far”. Models of estimated movement direction therefore included a fixed effect of movement endpoint distance relative to the straight line connection between home position and target, and models of estimated movement extent included a fixed effect of radial movement extent relative to the radial target distance. The coefficients of these fixed effects reflect spatial precision for estimated movement direction and extent, respectively (i.e., perceptual resolution of actual left/right and far/short differences). Our main hypothesis was that these effects of movement endpoint errors interact with the effect of condition. That is, spatial precision was expected to differ between conditions. The full model of estimated movement direction therefore included a fixed effect for movement endpoint distance relative to a straight line connection between home position and target, a fixed effect for condition, and a fixed effect for their interaction. Accordingly, the full model of estimated movement extent included a fixed effect for radial movement extent relative to target distance, a fixed effect for condition, and a fixed effect for their interaction. Random effects included random slopes per subject for each of these effects (Bates, Kliegl, Vasishth, & Baayen, 2015), and a random intercept per subject. In addition, we expected participants’ estimates to be biased by target locations. The full models of estimated movement direction and extent therefore also included a fixed effect for target coordinates in the x and y directions, respectively. Twelve quadrature points were used in the approximation of random effects. We report marginal coefficients.

To test each of the fixed effects, we compared the full model with a simpler model omitting the fixed effect to be tested. P-values for fixed effects were obtained by likelihood ratio tests (LRT; χ^2^ distribution). We also report model differences in the Bayesian Information Criterion (BIC) and the Akaike Information Criterion (AIC).

Plots were created in MATLAB, including functions pointSpread.m (Jonas, 2020) and error_ellipse.m (Johnson, 2020). For Figure 3E, 95% confidence interval ellipses were aligned in center coordinates and orientation, the latter based on eigenvectors of individual-subject covariance matrices.

## Results

We expected spatial estimates of movement endpoints to be more precise in the predictable as compared to the unpredictable condition. Furthermore, if movement corrections across trials rely on precise estimation of position, movement accuracy should be higher in the predictable condition.

For each experiment, we first report kinematics before turning to perceptual estimates of movement endpoints.

## Experiment 1

### Movement kinematics

**Figure 2A** depicts movement trajectories and movement speed of a typical subject in each condition. Across conditions, reaching movements were sufficiently accurate to cause the target to explode in 27.5 ± 7.8 % of trials (mean ± standard deviation (SD) across subjects), leaving considerable variability in movement endpoints around the target across the remaining trials (**Figure 2B**).

In both conditions, participants tended to stop too far to the left and to overshoot the target. Importantly, movement accuracy was significantly higher in the predictable compared to the unpredictable condition. The average distance between movement endpoint and target location was smaller in the predictable than the unpredictable condition (median: .74 cm vs. .86 cm (predictable and unpredictable, respectively), W=49.5, p=.007, matched rank biserial correlation (MRBC)=.64). Accordingly, the target exploded significantly more often in the predictable (31.5 ± 8.8%) compared to the unpredictable condition (23.5 ± 9.9 %, mean ± SD; t(22)=3.7, p=.001, d=.77; **Figures 2C** and **D**). Interestingly, peak movement velocity was higher in the predictable than the unpredictable condition (mean ± SD: 90.8 ± 18 cm/sec vs. 82.5 ± 11.7 cm/sec (predictable and unpredictable, respectively), t(22)=3.4, p=.003, d=.71).

### Perception of movement endpoints

**Figure 3A** depicts estimated movement endpoints, relative to actual endpoints, across all individuals. On average across conditions, participants misjudged the location of their movement endpoint by .81 cm (median across subjects). Participants tended to misjudge movement endpoints to the right and to underestimate movement extent, i.e., they reported movement endpoints to be closer to the target than they really were (compare Figures 3A and 2B).

Estimation accuracy was higher in the predictable as compared to the unpredictable condition. Movement endpoints were mislocalized, on average across individuals, by .7 cm in the predictable condition and by .88 cm in the unpredictable condition (W=47, p=.004, MRBC=.66). Accordingly, target explosion upon endpoint estimation occurred more frequently in the predictable than in the unpredictable condition (mean ± SD: 34.2 ± 11.5% (predictable) vs. 26 ± 10.1% (unpredictable), t(22)=3.78, p=.001, d=.788; **Figures 3B** and **C**). Importantly, endpoint estimations were also significantly more precise in the predictable than in the unpredictable condition, as expected. **Figure 3D** displays individual subjects’ 95% confidence ellipses of endpoint estimations, relative to actual endpoints. To facilitate comparison of ellipse sizes across conditions, **Figure 3E** shows individual subjects’ ellipses for each condition, centred and aligned in orientation. Variance of endpoint estimations was .55 cm in the predictable condition, and .84 cm in the unpredictable condition (median across subjects; W=43, p=.003, MRBC=.69). This difference persisted when taking into account differences in estimation accuracy. Average coefficients of variations in the two conditions were .88 ± .18 (predictable) and .97 ± .16 (unpredictable; mean ± SD; t(22)=2.32, p=.03, d=.485; **Figure 3F** and **G**).

Inter-individual differences in the precision of endpoint estimation correlated strongly with inter-individual differences in movement accuracy (predictable condition: R^2^=.86, p<.001; unpredictable condition: R^2^=.82, p<.001). This could suggest that perceptual precision enhances movement accuracy, e.g., by promoting larger trial-by-trial movement corrections. On the basis of Experiment 1 alone, however, we could not exclude the reverse, i.e., enhanced perceptual precision due to higher movement accuracy in the predictable condition. Endpoint estimation required moving a “copy” of the target from the target location to the estimated endpoint location using a trackball. In the unpredictable condition, the “copy” had to be moved, on average, further away from the target (Figures 2B and C). The cost of responding accurately was thus higher in the unpredictable condition. In addition, longer trackball movements in the unpredictable condition may have resulted in more (motor) noise at a response stage. To eliminate these potential confounds, we altered the method by which participants reported estimated endpoints in Experiment 2, while keeping all other aspects of the design identical.

## Experiment 2

### Movement kinematics

Experiment 2 replicated our finding that movement accuracy was significantly higher in the predictable compared to the unpredictable condition (**Figure 4A**). The average distance between movement endpoint and target location was smaller in the predictable than the unpredictable condition (mean ± SD: .9 ± .23 cm vs. 1.1 ± .36 cm (predictable and unpredictable, respectively), t(20)=2.79, p=.011, d=.61). Accordingly, the target exploded significantly more often in the predictable (25.7 ± 7.7%) compared to the unpredictable condition (19.6 ± 9.4 %, mean ± SD; t(20)=3.08, p=.006, d=.673; **Figures 4B** and **C**). Peak movement velocity was numerically higher in the predictable than the unpredictable condition (mean ± SD: 79.3 ± 21.7 cm/sec vs. 75.1 ± 17 cm/sec (predictable and unpredictable, respectively)), however, unlike in Experiment 1, this difference was not statistically significant (t(20)=1.13, p=.27).

### Perception of movement endpoints

As expected, we found that estimated endpoints relative to the target reflected actual movement error. Estimated movement direction (“left”/”right”) depended on the location of movement endpoints relative to a straight line connection between the home position and target (χ^2^(1)=29.4, p<.0001; .78 increase in log odds of responding “right” with each 1 cm increase in actual endpoint location, from left to right; ΔAIC=27.4, ΔBIC=32.4). Similarly, estimated movement extent (“too far”/”too short”) depended on the relative radial distances of actual movement endpoints and targets to the home position (χ^2^(1)=29.3, p<.0001; .37 increase in log odds of responding “too far” with each 1 cm increase in radial movement extent; ΔAIC=27.3, ΔBIC=26.2). For estimated movement extent, there was also a main effect of condition (χ^2^(1)=9.1, p=.003; .76 decrease in log odds of reporting “too far” in the predictable condition; ΔAIC=7.1, ΔBIC=6). Target coordinates along the x-axis significantly biased “left”/”right” responses (χ^2^(1)=42.6, p<.001; .76 increase in log odds of responding “right” with each 1 cm increase in the target’s x-coordinate towards the right; ΔAIC=40.6, ΔBIC=39.6), while target coordinates along the y-axis showed only a trend to bias “too far”/”too short” judgements (χ^2^(1)=3.3, p=.07;.2 increase in log odds of responding “too far” with each 1 cm increase in distance of the target from the home position; ΔAIC=1.3, ΔBIC=.3).

Importantly, there was a significant interaction between movement error and condition on estimated movement direction (χ^2^(1)=5, p=.026; ΔAIC=3, ΔBIC=2). The log odds of responding “right” as the actual movement endpoint moved 1 cm to the right were significantly higher in the predictable than in the unpredictable condition (by .6; **Figure 5A**). That is, participants had a spatially more resolved, i.e., more precise, perceptual representation of their actual movement endpoints. This interaction was in the same direction, but not significant, for estimated movement extent (χ^2^(1)=1.5, p=.23; ΔAIC=, 0.6, ΔBIC=1.6; **Figure 5B**).

## Discussion

Precise estimates of the location of body parts are important for movement planning and correction (Desmurget, Pelisson, Rossetti, & Prablanc, 1998; Desmurget, Rossetti, Jordan, Meckler, & Prablanc, 1997; but see Polit & Bizzi, 1978). The human position sense becomes more reliable when visual and proprioceptive information is integrated (van Beers et al., 1996, 2002). Often, however, movements unfold without visual feedback. Here, we show that, in the absence of visual feedback, human position sense relies partly on internal state estimation (Wolpert et al., 1995) derived from a predictive visuomotor model, which is tuned in both space and time. This internal estimation enhances precision of position sense, and improves movement accuracy.

On theoretical grounds, it has been proposed before that an internal model estimate contributes to perceptual precision in the context of movement. For example, Vaziri et al. (2006) argued that forward models may provide a separate source of information for perceptual inference, additional to sensory input. Following the idea of maximum-likelihood estimation as a principle of optimal integration (Ernst & Banks, 2002), the authors expected lower uncertainty of a final perceptual estimate when this internal model estimate is integrated with sensory input. To test this idea, Vaziri et al. (2006) turned to the oculomotor system and examined the variance with which healthy individuals localized a visual target from memory. In order to examine the effect of an internal model estimate on target localization, the authors compared a condition in which participants executed a saccade during the memory delay with a condition in which gaze remained static. It is well known that, before a saccade, receptive fields of neurons in several regions of the nervous system shift to the location in the visual field from which they will receive input after the saccade (e.g., Duhamel, Colby, & Goldberg, 1992; Merriam, Genovese, & Colby, 2003; Nakamura & Colby, 2002; Sommer & Wurtz, 2006). This shift has been taken as evidence of an internal model that estimates imminent eye movement based on an efference copy (Sommer & Wurtz, 2008). Following this idea, Vaziri et al. (2006) hypothesized that this internal estimate, available in the presence, but not absence, of a saccade, would decrease variance of target localization. Indeed, the authors found a reduction in variance in the saccade condition. However, the reason for this variance reduction in their study remains ambiguous. In the saccade condition, the visual target was presented foveally, i.e., with high visuospatial acuity, while it was only ever seen peripherally in the control condition, i.e., with lower spatial resolution. Higher visuospatial resolution at the time of encoding, rather than the presence of an internal model estimate, may thus explain why variance in target localization was lower in the saccade condition.

More recently, Yon et al. (2018) related perceptual precision to a predictive internal model in a functional magnetic resonance imaging (fMRI) study. They compared accuracy in decoding visual feedback of finger movements from fMRI signals while participants executed movements with the same finger as visually presented, or with another finger. They found higher decoding accuracy, and an fMRI signal suppression in voxels tuned away from the current visual stimulus, when participants moved the same finger as they concurrently saw on the screen. Yon et al. (2018) interpreted this as a sharpening of representations when sensorimotor predictions are met by sensory evidence. However, a different, possibly more parsimonious, account is to explain differences in fMRI decoding between these conditions through different degrees of cross-modal (proprioceptive vs. visual) sensory conflict, rather than assuming any role of a predictive model.

Finally, Bhanpuri et al. (2013) showed that an unpredictable resistive torque applied during movement decreases the precision of perceptual reports of movement extent when compared to a condition that does not involve any external torques. Bhanpuri et al. (2013) interpreted this as evidence that predictive models contribute to perceptual precision. However, applying resistive torques during movement impacts directly on sensory information in a spatial dimension that is highly task-relevant in their study, i.e., proprioceptive estimates of movement extent. Therefore, their investigation of perceptual precision in healthy individuals does not dissociate internal, i.e., predicted, vs. purely sensory spatial information (see below for their results from patients with cerebellar ataxia).

Our study minimizes ambiguities in previous studies. Unlike Yon et al. (2018), we operationalise predictability directly, via context, rather than via sensory congruency, avoiding potential confounds of cross-modal conflict. While Vaziri et al.’s (2006) and Bhanpuri et al.’s (2013) operationalisation of predictability influences task-relevant spatial information in sensory input, we hold spatial information constant across conditions and instead manipulate temporal stimulus predictability. This also allows us to show that spatial and temporal aspects of sensorimotor models are inherently coupled: Disrupting temporal predictability impairs spatial precision of perception (see also Bays et al., 2005; Blakemore et al., 1999; Kilteni et al., 2019).

Examining patients with cerebellar ataxia, Bhanpuri et al. (2013) demonstrated that active movement enhances precision of the perceived movement extent over passive movement only when functional integrity of the cerebellum is preserved. Because the cerebellum is considered critical for predictive state estimation (Miall, Christensen, Cain, & Stanley, 2007), they concluded that a cerebellum-dependent predictive state estimate contributes to perceptual precision. Our findings in healthy individuals point to a contribution of a specific, i.e., visuomotor, model to perceptual precision in the context of reaching. In addition, beyond perception, we demonstrate that movement accuracy is higher when a full, spatiotemporal, visuomotor model can be formed. One possible reason is that participants may correct future movements more efficiently when estimating movement endpoints with higher precision. Predictive models may thus influence movement in part via perceptual precision.

This has important potential implications for motor learning. Movement errors can arise from mis-planning despite unaltered sensorimotor contingencies, and then result in revised action selection, e.g., re-aiming (McDougle, Ivry, & Taylor, 2016). More precise estimation of movement endpoint, and, thus, movement error, likely facilitates re-aiming. On the other hand, movement error can also arise from a change in sensorimotor contingencies, and then result in adaptation of internal models (Shadmehr et al., 2010). A sudden change in sensorimotor contingencies is thought to lower precision of the present sensorimotor model (Tan, Wade, & Brown, 2016). Given our observation that internal models contribute to spatial precision of position sense, the latter should thus become temporarily less precise upon a change in sensorimotor contingencies. This could explain why re-aiming through revised action selection, which likely depends on precise estimation of movement error, lags internal model adaptation (Mazzoni & Krakauer, 2006; Taylor & Ivry, 2011), an idea we are currently testing.

Two aspects of our experimental manipulation could have caused the observed decrease in perceptual precision in the unpredictable condition. The onset time of visual feedback in the unpredictable condition was both jittered and delayed. The jitter made visual feedback unpredictable. However, the delay, too, may have contributed to lowered predictability. Delayed feedback results in reduced internal model calibration in visuomotor adaptation paradigms, as reflected in diminished aftereffects of reaching (Brudner, Ivry, Taylor, Graeupner, & Kethidi, 2016; Schween & Hegele, 2017). Whether the jitter, the delay, or both, reduced the extent to which participants based their perceptual estimates on a visuomotor model remains to be tested.

Besides a visuomotor model estimation, participants could have based perceptual reports on proprioception. However, the spatial pattern of variance in our study contrasts with the typical spatial pattern of variance of proprioception. Across the two experiments, estimation variance was larger in depth (movement extent) than in azimuth (movement direction; Figures 3A, 3D, and 5). Proprioceptive estimates, on the other hand, typically show the opposite pattern, i.e., larger variance in azimuth than in depth (when the arm is extended; van Beers et al., 2002; van Beers, Sittig, & Denier van der Gon, 1998). The observed spatial pattern of variance in endpoint estimation thus speaks against a dominant role of proprioception in our study.

The spatial pattern of variance observed in our study may instead be explained by spatially dependent uncertainty in internal model estimates. It is reasonable to assume that forward model estimation is less precise in depth than in azimuth, given that movement variability is typically larger in movement extent than in movement direction (e.g., He et al., 2016; van Beers, 2003). In addition, our finding that peak velocity was higher in the predictable than in the unpredictable condition, at least in Experiment 1, points to a stronger reliance on feedforward control mechanisms in the predictable condition.

To conclude, we show that a spatiotemporal, visuomotor model enhances the precision of human position sense, and thereby increases movement accuracy. A critical aspect of quality of perception – its certainty, and reliability – is thus influenced not only by the quality of sensory input, but also by predictive control mechanisms of motor output. This could be particularly relevant in the context of active sensing, and possibly explain interactions of different, and sometimes opposing, motor learning mechanisms.

## Acknowledgements

We thank Patrick Riese for excellent technical support. We thank all participants for their time and effort, and all members of the Motor Learning research group for their valuable feedback on the manuscript.

## Competing interests

The authors report no conflicts of interest.

## Funding

M.-P. Stenner was supported by a VolkswagenStiftung Freigeist Fellowship, AZ 92 977, and received funding from a Deutsche Forschungsgemeinschaft Sonderforschungsbereich Grant, SFB-779, TPA03.

## Notes

### Competing Interest Statement

The authors have declared no competing interest.

